# CRISPR/Cas9 screening using unique molecular identifiers

**DOI:** 10.1101/114355

**Authors:** Bernhard Schmierer, Sandeep K. Botla, Jilin Zhang, Mikko Turunen, Teemu Kivioja, Jussi Taipale

## Abstract

Loss of function screening by CRISPR/Cas9 gene knockout with pooled, lentiviral guide libraries is a widely applicable method for systematic identification of genes contributing to diverse cellular phenotypes. Here, random sequence labels (RSLs) are incorporated into the guide library, which act as unique molecular identifiers (UMIs) to allow massively parallel lineage tracing and lineage dropout screening. RSLs greatly improve the reproducibility of results by increasing both the precision and the accuracy of screens. They reduce the number of cells and sequencing reads needed to reach a set statistical power, or allow a more robust screen using the same number of cells.

---

Pooled CRISPR/Cas9 loss of function screening is a powerful approach to identify genes contributing to a wide range of phenotypes^1^. A library of guide sequences is integrated lentivirally into Cas9-expressing cells, which are then subjected to a selection pressure. Relative guide frequencies in the population before and after selection are quantified by next generation sequencing (NGS) to determine depleted and enriched guides.

The approach has been applied successfully^2–5^, but suffers from several shortcomings: First, the presence of a guide does not necessarily cause loss of the corresponding gene, and cells sharing the same guide have distinct genotypes and phenotypes. Second, identification of guides that are under negative selection can be confounded by random drift and undersampling. Third, growth characteristics of individual cells can vary substantially^6, 7^ and the site of viral integration can affect the phenotype. For these reasons, each guide needs to be present in a large number of cells. In conventional screens, only the sum of all cells with a specific guide is measured, and no information regarding the distribution of cell behaviors can be obtained. Optimal identification of hit genes would require a method that individually tracks clonal lineages derived from single virus-transduced cells.

Here, we address these issues by incorporating an RSL into the guide-library plasmid (**Fig. 1a**) to allow tracing of thousands of individual virus-transduced cell lineages in a CRISPR screen. Depending on the kinetics of editing, these unique molecular identifiers (UMIs)^8^ either trace single clones of identically edited cells, or small populations of sublineages with different editing outcomes at the same locus (**Supplementary Fig. 1a**). Such massively parallel lineage tracing enables both lineage dropout analysis (LDA), and the creation and analysis of internal replicates (IRA), while retaining the option of conventional, total read count analysis (TCA, **Fig. 1b**). Perhaps counterintuitively, analysis of hundreds to thousands of RSL-labelled cell lineages per guide neither requires more cells per guide, nor deeper sequencing. The RSL approach simply splits the total guide read count obtained to read counts representing individual constituent cell lineages, thus increasing the amount of information that is obtained, and consequently improving both precision and accuracy of the screen.

To demonstrate the power and flexibility of the approach, we screened the human colorectal carcinoma cell line RKO for essential genes with an RSL-guide library targeting 2325 genes with 10 guides per gene^5^. For experimental details see **Supplementary Information**. Briefly, Cas9 expressing RKO cells were transduced with the guide library, and samples were taken at Day 4 and Day 28 after transduction (control and treatment time points, respectively). Guide frequencies in the two samples were then assessed by NGS. The experiments were run at far larger screen size (we define ‘screen size” as the number of cells per guide sequence) and sequencing depth (reads per guide) than previous screens ^4, 5^. Such redundancy allows subsequent subsampling using RSL information, and robust testing of different analytical methods at varying screen sizes (**Fig. 1c**).

The plasmid library input contained 78 million unique RSL-guide combinations, 93% of which were also detected in the virus-transduced cell populations (**Supplementary Fig. 1b**). Based on the Poisson-distribution, this indicates that about half of the RSL-guides were incorporated into one or two cell lineages. Because only a subset of the cells can be harvested at each time-point, undersampling is unavoidable, and some RSL-guides and thus cell lineages were present only in one of the time points (**Supplementary Fig. 1b, right**). Such undersampling and loss of cell lineages occurs whether or not RSLs are present, however goes undetected in their absence. With RSLs, the effect becomes apparent and can be used in quality control of individual experiments as well as in filtering out inconsistently sampled lineages prior to data analysis.

RSL-labelled, distinguishable guide sequences can be used to split the data into internal replicates, which in turn allow the usage of classical statistical tools to test for significant differences. To demonstrate the approach, RSL-guides were binned into 64 internal replicates per guide. The median effect size (**Fig. 1d**) as well as a median-based version of strictly standardized mean difference (SSMD)**^9^** were then used to rank the guides (internal replicate analysis using SSMD, IRA/SSMD, **Supplementary Fig. 1c**). In addition, RSL-labelled guides enable lineage dropout screening, where gene hits are called solely based on the number of lost RSL-guide lineages (lineage dropout analysis, LDA, **Fig. 1e**).

To evaluate IRA/SSMD and LDA, and to compare them with conventional TCA performed with the pipeline MAGeCK^10^, we assessed the ranks of a set of known essential genes (accuracy), and the hit gene overlap between experimental replicates (precision). In principle, RSL-based methods should outperform TCA when the number of cells per guide is relatively low, and their benefit should progressively decrease as the number of cells per guide approaches infinity. Thus, the comparisons were performed using the complete dataset, and subsamples of the data that were similar in sample size to published screens ^4, 5^ (**Fig. 2**).

Both IRA/SSMD and LDA were more accurate than TCA, as indicated by lower hit ranks of 20 known essential, ribosomal proteins (**Fig. 2a, Supplementary Fig. 1d**). Both IRA/SSMD and LDA were also more precise than TCA, with much improved replicate concordance between the top-ranked 5% of genes (**Fig. 2b**). Consistently with the theoretical considerations, our analysis revealed that the RSL-based methods were far more robust at smaller screen sizes than TCA. Even when the screen size was approximately one order of magnitude larger, TCA was still inferior to either RSL-based method. At smaller screen size, the number of highly significant hit genes (FDR < 1%) was massively increased in lineage dropout analysis when compared to total readcount analysis (**Fig. 2c**).

To summarize, RSLs dramatically improve accuracy, precision, and statistical power in CRISPR/Cas9 screening. The RSL strategy is not limited to CRISPR knockout screening, but can be applied in other screening methods such as CRISPR-dependent inhibition or activation screens^2, 11^. We expect the RSL method to become instrumental in the interrogation of small genomic features, e.g. exons, promoters, and even individual transcription factor binding sites. In many of these cases there is just one possible guide sequence, and the inclusion of RSLs is the only way to obtain the replicates that are required for hit calling. In the absence of precise knowledge of both on-and off-target activity, inclusion of multiple guide positions is however still important, and rescue experiments and/or analysis of the mutational spectrum of the cutsite are necessary to establish that the mutation induced by the guide results in the observed phenotype. Incorporation of RSLs is technically straightforward, and does not require a higher number of cells or sequencing reads compared to conventional approaches. In contrast, RSLs give the same statistical power at lower number of cells and/or sequence reads, improving the economy of CRISPR/Cas9 screens. Conversely, RSLs improve accuracy and precision at a given number of cells per guide, which is particularly advantageous in cases where cell numbers are limiting, such as in primary cells.

**Figure 1.**
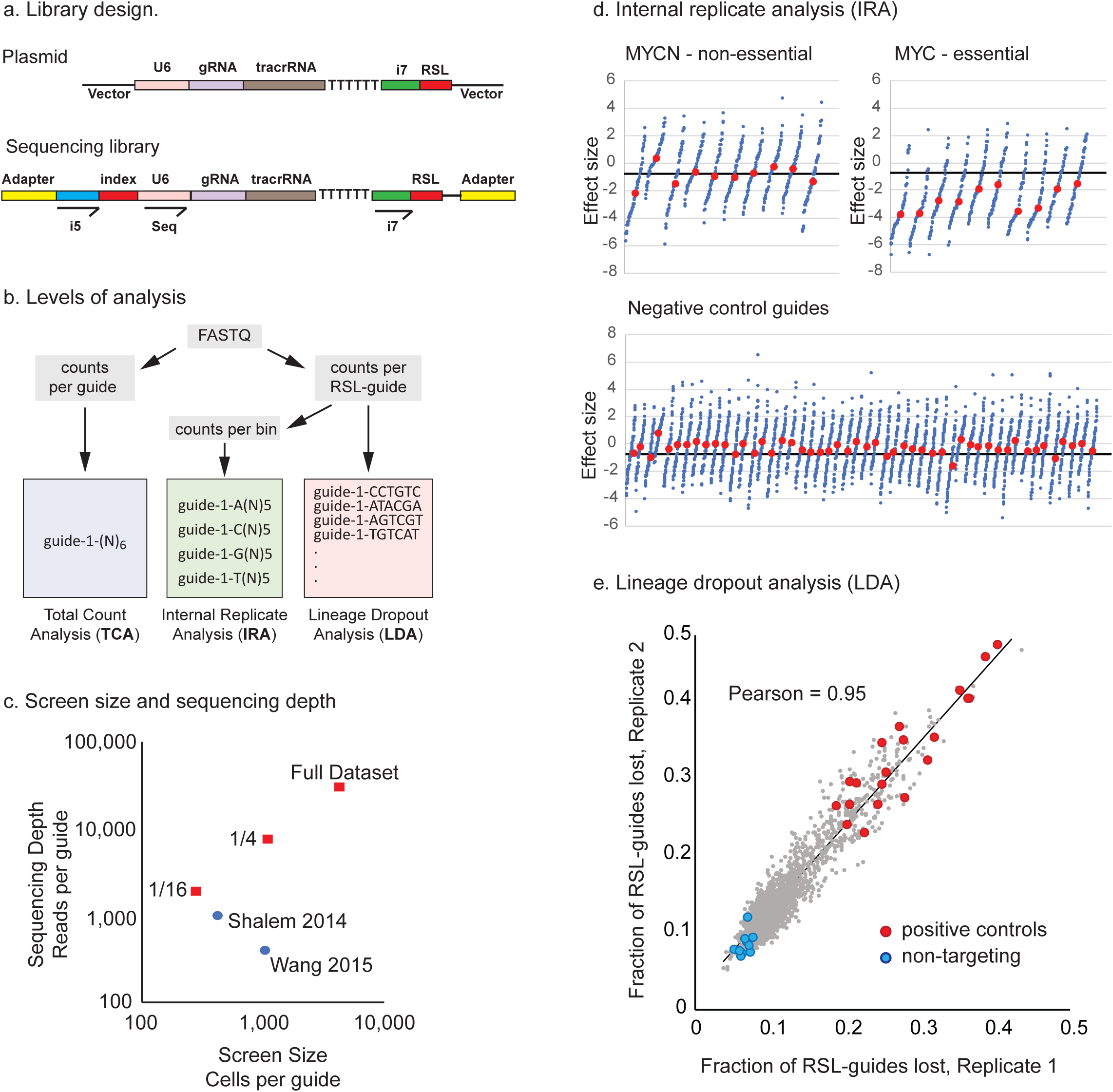
**a. Library design. Top.** Guide plasmid. The plasmid library contains an i7 index read primer and an inert, untranscribed RSL downstream of the tracrRNA. **Bottom.** Sequencing library. Sequencing was done with a custom primer (Seq) placed directly upstream of the guide (gRNA). The sample index and RSL were read as two index reads with illumina i5 and i7 index primers, respectively (20+6+6 sequencing cycles). **b. RSL guides allow additional methods of analysis.** In Total guide read Count Analysis (TCA, left) RSL information is ignored and only the sum of readcounts for all RSL-guides is taken into account. In internal replicate analysis (IRA, middle), readcounts of RSL-guides are binned such that internal replicates are created for each guide. The example shown bins into four internal replicates. In lineage dropout analysis (LDA, right) each RSL-guide is monitored separately. **c. Screen size and sequencing depth.** The screens were performed at a very large screen size of roughly 4500 cells per guide and sequenced to a depth of 30,000 reads per guide. Using RSL information, the data from these oversized experiments were then subsampled bioinformatically to approximately one quarter and one sixteenth, to test different analysis methods at different screen sizes. The corresponding values for two published screens are indicated for comparison ^4, 5^ **d. Internal replicate analysis (IRA).** RSL-guides were binned to create 64 internal replicates. Effect sizes (log2 fold change in readcount between Day 4 and Day 28 after virus transduction) for each bin are plotted in ascending order, 10 guides each for MYCN (**top left**) and MYC (**top right**), as well ass 50 representative non-targeting guides (**bottom**, these non-cutters seem to have a small fitness advantage). Red dots, median effect size (MES) of the 64 internal replicates; black line, MES of all guides in the library. Hits for this type of data were called from MES and SSMD (**Supplementary Figure 1b,** see Supplementary Information for details). **e. Lineage dropout analysis (LDA).** The average fraction of RSL-guides lost from day 4 to day 28 in each experimental replicate is plotted for each gene. Red, positive controls; blue non-targeting controls; black line, linear regression. The number of virus-transduced cell lineages lost is the most direct readout of the guide effect on cell viability.

**Figure 2.**
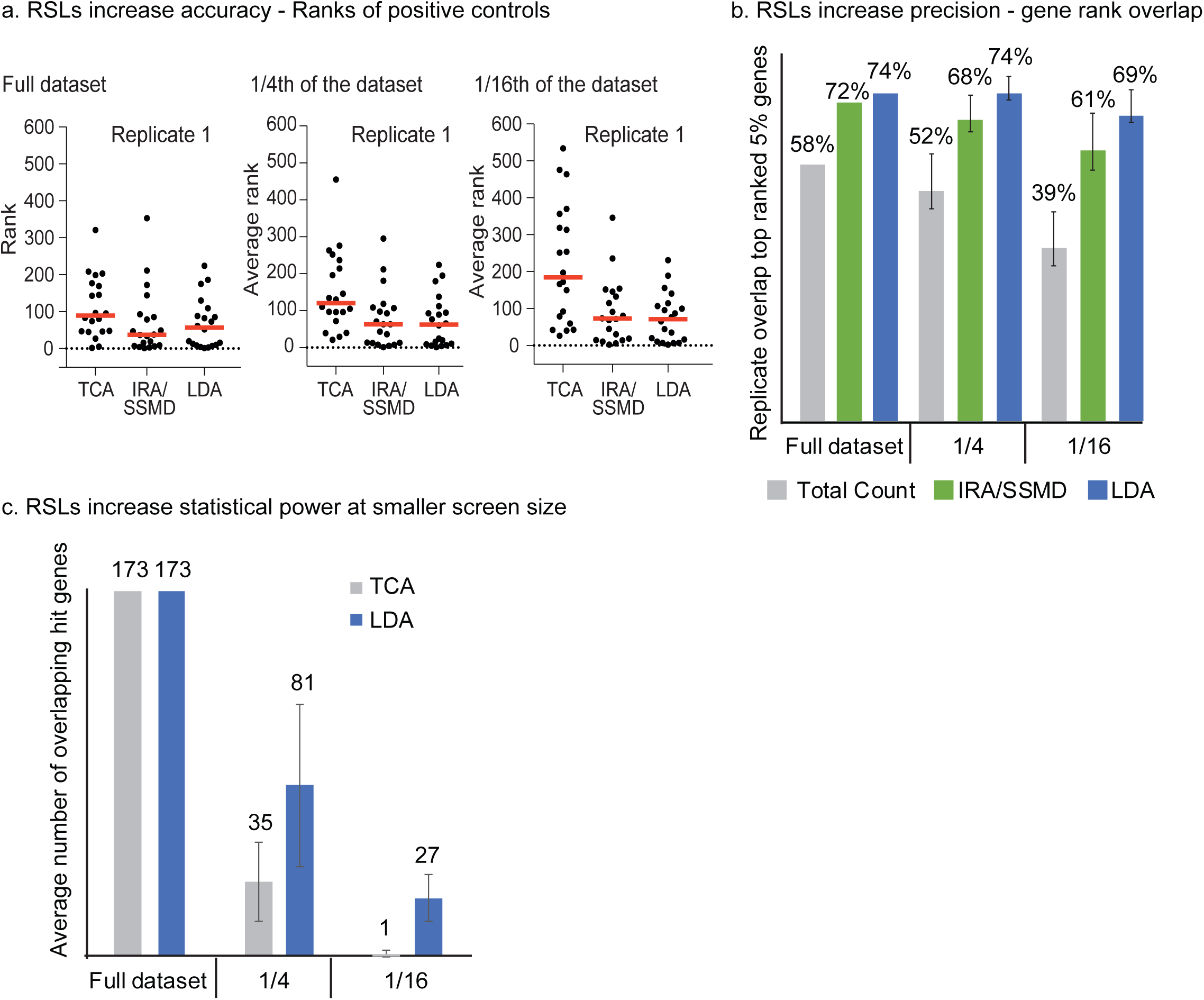
**a. RSLs increase accuracy of hit calling.** Ranks of known positive controls (20 ribosomal proteins out of a total of 2,335 interrogated genes) in one experimental replicate for the full screen size (left), as well as one quarter (middle) and 1/16 (right) of the full screen size. Red line, median rank. At all screen sizes, IR/SSMD analysis and LDA assigned lower ranks to the positive controls than TCA. The variance of the ranking increased substantially with decreasing screen size in TCA, but not in the two RSL-based methods. **b. RSLs increase the precision of gene ranking.** Average percent overlap of the top-ranked 5% of genes (116 genes) between two experimental replicates. Error bars, standard deviation of four subsamples. LDA is the most precise method, followed by IRA/SSMD. Again, both RSL-based methods are superior to TCA and much more robust at smaller screen sizes. **c. RSLs boost statistical power at small screen size.** Hit gene overlap between experimental replicates (<1% false discovery rate) at full screen size, one quarter and one sixteenth of the full screen size. Error bars, standard deviation for hit gene overlap between four subsamples in experimental replicate 1 and four subsamples in experimental replicate 2 (16 comparisons in total). Only at full screen size, TCA matches LDA. At more practical screen sizes, LDA has much higher statistical power and identifies considerably more hit genes.

## Accession codes

Read data: European Nucleotide Archive, PRJEB18436. Scripts will be made available on Github under public license.

## Acknowledgements

The authors would like to thank Drs. Inderpreet Kaur Sur, Jenna Persson and Minna Taipale for suggestions on the manuscript. Part of this work was carried out at Karolinska High Throughput Center (KHTC) and the High Throughput Genome Engineering Facility (HTGE) funded by SciLifeLab.

## Authorship contributions

B.S., S.K.B. and J.T. developed the approach, B.S., S.K.B. and M.T. performed the experiments, B.S., S.K.B, J.Z. and T.K. analyzed the data, B.S. and J.T. wrote the manuscript.

## Schmierer et al., Supplementary Figure 1

**Supplementary Figure S1.**
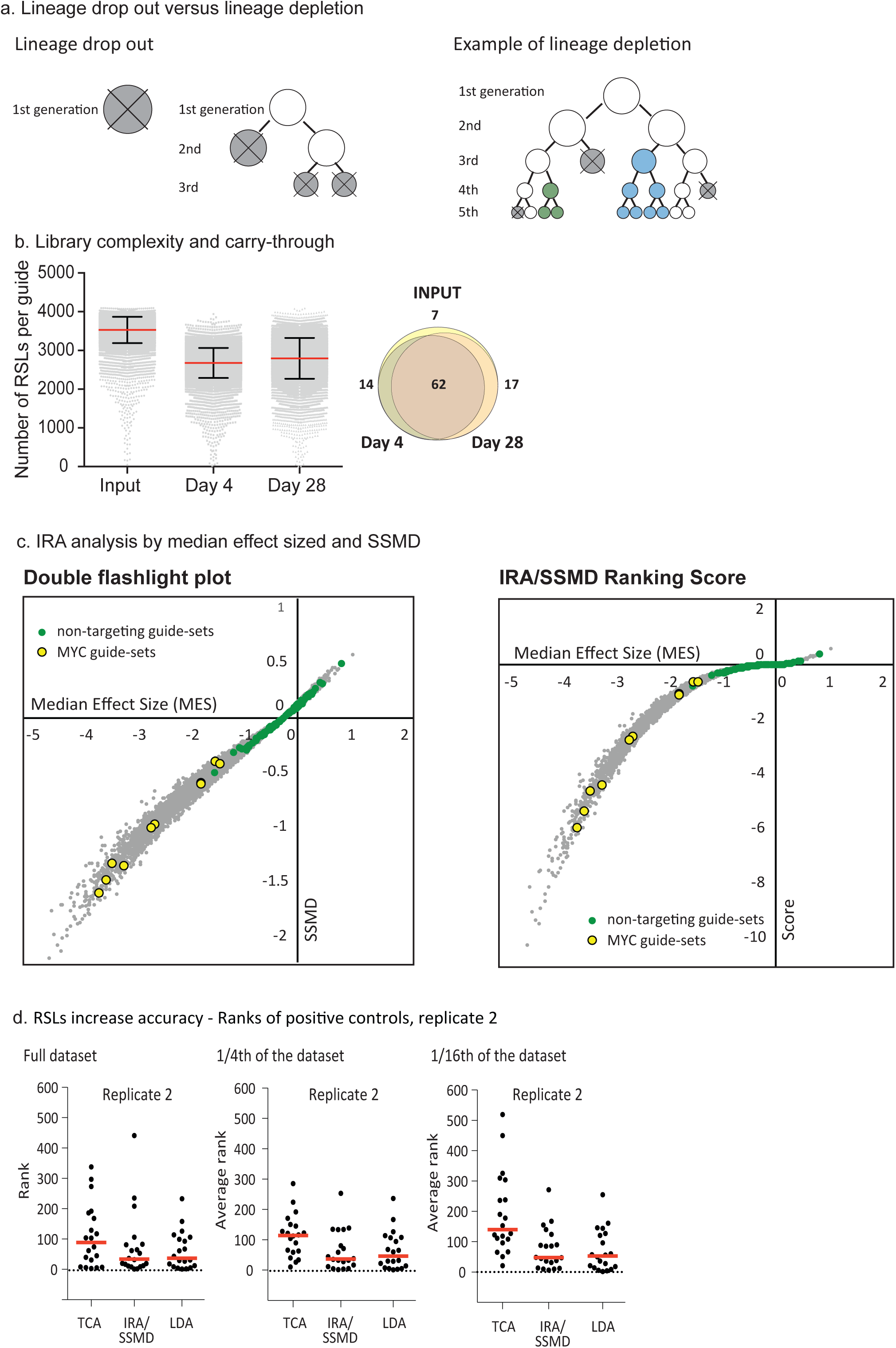
**a.** Lineage dropout versus lineage depletion. Depending on the kinetics of editing, single cell lineages harboring a single RSL-guide against an essential gene can either disappear (drop out) or decrease in their abundancy (depletion). **Left.** Dropout happens if the editing occurs early on, either before the cell can divide, or in several independent events at later time points (grey, dead cell; white, unedited cell). **Right**. In lineage depletion, editing occurs either after several cell divisions and/or with several different outcomes, some of which will retain gene function of the essential gene (blue and green edits). The traced lineage is then comprised of several sublineages. **b. Number of distinct sequences carried through the experiment in one representative experimental replicate. Left.** Boxplot. An average of 3600 RSLs per guide in the plasmid library reduced to an average of 2800 RSLs per guide in the samples taken from the cell populations. Many of these RSLs have very low read counts, those were filtered out and not used for downstream data analysis. **Right.** Venn diagram. Both timepoints together covered 93% of input RSL-guides (78 million unique sequences in the cell population). The overlap between day 4 and day 28 was two thirds, with about one sixth of sequences found either only in Day 4 or only in Day 28. This is a consequence of unavoidable undersampling, which also occurs in the absence of RSLs, however in their presence becomes apparent and allows removal of inconsistently sampled lineages. **c. Internal replicate analysis by strictly standardized mean difference (IRA/SSMD). Left.** Double flash-light plot. SSMDs for all >23,000 guide sets are plotted against their median effect sizes (MES). Green circles indicate non-targeting control guide sets, yellow circles indicate the 10 guide-sets targeting MYC. **Right.** Ranking score. A hit score was defined as the product of MES and SSMD. This score is negative for depleted guides and positive for enriched guides, and was used for guide ranking. The score was plotted against the median effect size for all guides. Green circles indicate non-targeting control guide sets, yellow circles indicate the 10 guide-sets targeting MYC. **d. RSLs increase accuracy of hit calling.** As Fig. 2a, but for the second experimental replicate.

### Supplementary Information

**Oligo synthesis and library cloning.** The guide library targeted 2325 genes and contains a total of 23,279 guides. The targeted gene set contains all human transcription factors ^1^, other genes of interest as well as ribosomal proteins as positive controls and 101 non-targeting guides as negative controls. All sgRNA sequences used in this library were taken from a previously published, genome-wide library^2^ (Supplementary file *RSL_guide_library.csv*). The 5’ part of the library construct (blue + black, 122bp), containing the sgRNA was synthesized by oligo array (CustomArray). The 3’ part containing the RSL and the Illumina i7 index primer sequence (green) was synthesized as a single 119 bp oligo (black+green+red). These two oligos were annealed to each other at the overlapping part (black) and double stranded by PCR (T_M_ = 64C) using outer primers (underlined).

~~~
GGCTTTATATATCTTGTGGAAAGGACGAAACACCGNNNNNNNNNNNNNNNNNNNNgtttAagagctag
~~~

~~~
aaatagcaagttTaaataaggctagtccgttatcaacttgaaaaagtggcaccgagtcggtgcTTTTT
~~~

~~~
TgatcggaagagcacacgtctgaactccagtcacNNNNNNaagcttggcgtaactagatcttgagaca aa
~~~

The PCR product was cloned by Gibson assembly into the lentiviral vector pLenti-Puro-AU-flip-3xBsMBI, which was created by modifying lentiGuide-Puro (a gift from Feng Zhang, Addgene #52963) by replacing the sequence

~~~
gttttagagctagaaatagcaagttaaaataaggctagtccgttatcaacttgaaaaagtggcaccga gtcggtgcTTTTTT
~~~

with

~~~
gtttAagagctagaaatagcaagttTaaataaggctagtccgttatcaacttgaaaaagtggcaccga gtcggtgcTTTTTTcgtctct).
~~~

#### Gibson assembly, transformation and amplification of the library

100 ng vector and 12 ng insert where assembled in a total reaction volume of 100 μl (NEBuilder^®^ HiFi DNA Assembly Master Mix, NEB). The reaction was cleaned via a Minelute reaction cleanup column (Qiagen) and transformed into 6 x 50 μl electrocompetent *E. coli* (Endura(™) ElectroCompetent Cells, Lucigen) using a 1.0 mm cuvette, 25μF, 400 Ohms, 1800 Volts. Bacteria were plated on several 24x24 cm agar plates and colonies were grown overnight. Colonies were scraped into LB medium and the contained plasmids were isolated by Maxiprep.

#### Library packaging

The library was packaged in HEK 293T cells by cotransfecting the library plasmid and the two packaging plasmids psPAX2 (a gift from Didier Trono, Addgene #12260) and pCMV-VSV-G (a gift from Bob Weinberg, Addgene # 8454) in equimolar ratios. After 48 hours, the virus-containing supernatant was concentrated 40-fold using Lenti-X concentrator (Clontech), aliquoted for one time use and stored at -140C.

#### Cell lines and cell culture

All the cells used in this study were purchased directly from ATCC. Cells were regularly tested for mycoplasma using the Mycoalert detection kit (Lonza; cat# LT07-218).

#### Creating editing-proficient Cas9 cell lines

To rapidly generate editing-proficient cell lines, we synthesized a lentiviral construct (pLenti-Cas9-sgHPRT1) that encodes a codon optimized WT-SpCas9 that is flanked by two nuclear localization signals (derived from lenti-dCAS-VP64_Blast, a gift from Feng Zhang, Addgene #61425). In addition, the construct codes for blasticidin resistance, and carries an sgRNA against HPRT1 

~~~
(GATGTGATGAAGGAGATGGG).
~~~

 HPRT1 loss confers resistance to the antimetabolite 6-thioguanine (6-TG). Lentivirally transduced cells were selected in 5 μg/ml Blasticidin and after one week to 10 days additionally with 5μg/ml 6-TG until control cells had died. Only cells that both express Cas9 and are editing proficient, as indicated by loss of HPRT1 function, will survive. The method allows rapid establishment of a pool of editing proficient cells. Compared to single cell clones, this method retains the genetic heterogeneity of the original cell line, avoids potential clonal effects of the particular integration site of Cas9, and greatly accelerates cell line generation. These benefits need to be weighed carefully against possible disadvantages, such as synthetic lethality with HPRT1 loss, or potential effects of the presence of a second guide in the cell.

#### Library transduction

Per replicate, 100 million RKO Cas9 cells were transduced with the library virus. Cells were then selected for guide integration and expression by 1 μg/ml puromycin selection for 48 hours. A proportion of cells will contain more than one guide. Because of the vast number of RSL-guides, any ineffective passenger guides will associate with effective guides randomly and will not be significantly enriched or depleted in the population.

#### Cell propagation and sample preparation

Cells were kept in culture for a total of 28 days after transduction by sub-culturing them every three to four days. 100 million cells were reseeded at each split, and genomic DNA was prepared from 50 – 80 million cells at Days 4 and 28 after transduction. Day 4 after transduction was considered the control time point.

#### Preparation of the sequencing library from genomic DNA

The sequencing library preparation consists of 3 PCR steps, PCR1 amplifies the genomic region containing the guide sequence using the primers 1F and 1R. PCR2 and PCR3 then incorporate the Illumina adaptors with primers 2F/2R and 3F/3R, respectively. 3F contains the Illumina index for multiplexing, indicated by NNNNNN in the sequence given.

Genomic DNA was isolated using Blood and Tissue Maxi Kit (Qiagen), and 200 μg, theoretically corresponding to 30 million diploid cells, were used as PCR template in 40 parallel PCR reactions (5 μg template DNA each) using KAPA HiFi HotStart polymerase (KAPA Biosystems). After 14 cycles, the reactions were pooled. PCR2 used 5 μl of pooled PCR1 as template and was run for 19 cycles, PCR3 used 2μl of PCR2 as template and was run for 14 cycles. The resulting product of 288 bp was gel purified and sequenced on an Illumina HiSeq 4000 instrument using single read 20 cycles plus two 6 bp index reads, where index read 1 reads the RSL and index read 2 reads the Illumina sample index.

1fw GGACTATCATATGCTTACCGTAACTTGAAAGTATTTCG
1ref CTTTAGTTTGTATGTCTGTTGCTATTATGTCTACTATTCTTTCC
2fw TCTTTCCCTACACGACGCTCTTCCGATCtcttgtggaaaggacgaaacac 2rev AGAAGACGGCATACGAGATctgccatttgtctcaagatctagttac
3fw AATGATACGGCGACCACCGAGATCTACAC NNNNNN TCTTTCCCTACACGACGCTCTTCCG
3rev CAAGCagaagacggcatacgagatctgccatttg

The final library product was sequenced with a custom primer and the i5 and i7 index primers (underlined) by running 20+6+6 cycles on the Illumina HiSeq4000.

~~~
AATGATACGGCGACCACCGAGATCTACAC[i5]**NNNNNN**TCTTTCCCTACACGACGCTCTTCCGATCt
~~~

~~~
cttgtggaaaggacgaaacacCG**NNNNNNNNNNNNNNNNNNNN**gtttAagagctagaaatagcaagttTaaataaGgctagtccgttatcaacttgaaaaagtggcaccgagtcggtgcTTTTTTgatcggaagag cacacgtctgaactccagtcac[i7]**BBBBBB**aagcttggcgtaactagatcttgagacaaatggcag ATCTCGTATGCCGTCTTCTGCTTG
~~~

#### Scripts used for counting RSL-guides and for binning

RSL-guides were counted in the original fastq files with the Perl scripts *BatchRun-pub2.pl*, which requires the script *GuideUMI_pub2.pl*. Binning of RSL-guide counts was done using the script *countTruncatedRSLs.pl*. Generally, sequences whose total readcount in control and treatment was less than five were filtered out prior to data analysis.

### SSMD analysis of read count data

#### Normalization

In RNASeq, methods such as median normalization are commonly preferred to total read-count normalization, mainly to compensate for the effect of a few very highly expressed genes that can take up a significant proportion of the total read count. CRISPR/Cas9 screening data are comparably well balanced and we thus chose the most basic normalisation method, total read count normalisation, to compensate for different sequencing depths. *c*_*ij*_ and *t*_*ij*_ represent the raw read counts for RSL-guide *j* in guide-set *i* for control (Day 4 after lentiviral transduction) and treatment (Day 28 after lentiviral transduction), respectively. The normalised read counts 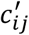 and 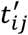 are then

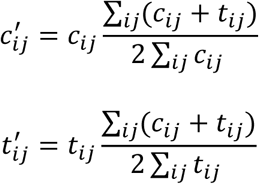

#### Median effect size and variability of the guide-sets

We defined the effect size *ES*_*ij*_ for each RSL-guide *i* in guide-set *i* as the log2 of the fold change between treatment count and control count. To handle total loss of an RSL-guide in the treatment sample, we added a pseudo-count of 1 to all counts:

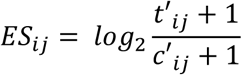

Next, we calculated the median effect size for guide set *i*, *MES*_*i*_, and the median of the absolute deviations (MAD) of all RSL-guides *j* in guide-set *i* from *MES*_*i*_

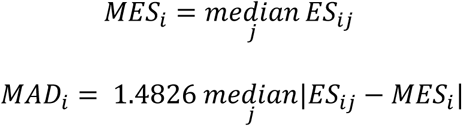

The factor 1.4826 was chosen such that the MAD is approximately equal to the standard deviation under the assumption of normal distribution^3^.

#### Median effect size and variability of the control guide-sets

The RSL library contains 101 non-targeting guide-sets. We calculate a single median effect size and MAD for this control set in the following way:

Median effect size of all non-targeting RSL-guides

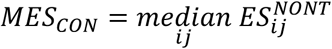

Median absolute deviation of all non-targeting RSL-guides:

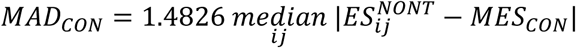

#### Strictly standardized mean difference (SSMD)

SSMD is a measure for the significance of the difference in behaviour of sample *i* and the non-targeting controls. It takes into account both the effect size and the variability of the data.

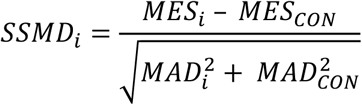

For samples with relatively small effect size, the SSMD can still become large if the spread is small. We thus introduce a score in which the effect size weighs more strongly, and which is used as a ranking parameter:

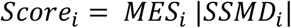

For hit calling, the average score and standard deviation were calculated for all non-targeting guide sets. The script used in these calculations is *SSMD.sh*, which calls the script R-script *SSMD.R*. Guide-sets were then ranked according to their score and the resulting ranked list was analysed with α-RRA, a robust rank aggregation algorithm as implemented in the ’pathway” function of MAGeCK^4, 5^ using Supplementary file *RSL_guide_library.gmt.*

### Lineage dropout

An RSL-guide was considered a dropout if it had less than two readcounts in the treatment time point. The numbers of RSLs per guide at Day 4 and Day 28 were then used as input in MAGeCK to obtain a ranked gene list and FDRs.

### Subsampling

For subsampling the full data set, RSL-guides were grouped according to their RSL-sequence. For medium screen size, the whole dataset was split into four groups (RSLs starting with A,C,G and T). For small screen size, the whole dataset was split into 16 groups, the first four of which (AA,AC,AG,AT) were used for analysis.

